# Nigratine as first-in-class dual inhibitor of necroptosis and ferroptosis regulated cell death

**DOI:** 10.1101/2020.12.15.422885

**Authors:** Claire Delehouzé, Arnaud Comte, Marcelle Hauteville, Peter Goekjian, Marie-Thérèse Dimanche-Boitrel, Morgane Rousselot, Stéphane Bach

**Affiliations:** SeaBeLife Biotech, Place Georges Teissier, 29680, Roscoff, France; Sorbonne Université, CNRS, UMR8227, Integrative Biology of Marine Models Laboratory (LBI2M), Station Biologique de Roscoff, 29680 Roscoff, France; Université de Lyon, CNRS UMR 5246, ICBMS, Chimiothèque, Université Claude Bernard Lyon 1, F-69622 Villeurbanne, France; Laboratoire de Biochimie Analytique et Synthèse Bioorganique, Université de Lyon, Université Claude Bernard Lyon 1, F-69622 Villeurbanne, France; Université de Lyon, CNRS UMR 5246, ICBMS, Laboratoire Chimie Organique 2- Glycosciences, Université Claude Bernard Lyon 1, F-69622 Villeurbanne, France; INSERM UMR 1085, Institut de Recherche sur la Santé, l’Environnement et le Travail (IRSET), F-35043 Rennes, France; Biosit UMS 3080, Université de Rennes 1, F-35043 Rennes, France; Sorbonne Université, CNRS, FR2424, Plateforme de criblage KISSf (Kinase Inhibitor Specialized Screening facility), Station Biologique de Roscoff, 29680 Roscoff Cedex, France

**Keywords:** dual-target inhibition, regulated necrosis, polypharmacology, ferroptosis, necroptosis

## Abstract

Nigratine (also known as 6E11), a natural flavanone derivative, was characterized as highly specific non-ATP competitive inhibitor of RIPK1 kinase, one of the key component of necroptotic cell death signaling. We show here that nigratine inhibited both necroptosis (induced by Tumor Necrosis Factor-α) and ferroptosis (induced by glutamate, erastin or RSL3 small chemical compounds) with EC_50_ in the µM range. Altogether, the data obtained showed that nigratine is the first-in-class dual inhibitor of necroptosis and ferroptosis cell death routes and opened new therapeutic avenues for treating complex necrosis-related diseases.

## INTRODUCTION

Advances in systems biology have revealed that single-target compounds are less efficient in preventing or cure complex diseases such as neurodegenerative diseases^1,2^. Drug discovery failures in complex cases, including Alzheimer’s disease (AD), may suggest that the definition of a “magic bullet” (drug selective for a single molecular target), the scientific concept developed by Paul Ehrlich more than one century ago^3^, should be tempered. The development of multi-target-directed ligands (MTDLs) for AD treatment is a notable example of promising therapeutic strategies^2^. Moreover, the most clinically effective central nervous system (CNS) drugs -such as clozapine for treatment of schizophrenia-act as “magic shotguns”: a non-selective drugs with pleiotypic actions^4^. These compounds are effective in treating complex human disorders because they are able to modulate the multiple targets involved in the pathophysiological processes, a strategy called polypharmacology. Synergistic effects are also among the main advantages of this combined therapeutic approach^5^. This strategy is now widely used in drug development as, from 2015 to 2017, 21% of the new molecular entities (NMEs) approved by the food and drug administration (FDA) are multi-target drugs (34% were single-target small molecules)^6^.

The systemic breakdown of physiological networks is not only restricted to the brain tissue and was described in numerous others including kidneys and liver. Functionally redundancy is typical features of such diseased networks and is well described for regulated cell death (RCD) signaling^7^. Indeed, this redundancy may reflect a particular evolutionary history for cell suicide and autophagic, apoptotic or necrotic elements might have been added to an ancestral death mechanism. Ancestral cell death routes probably include ferroptosis, a non-apoptotic cell death that is catalyzed by iron^8^, shown to be functional in cyanobacteria, components of phytoplankton communities evolving over 2.7 billions years ago^9^. As mentioned by Golstein and Kroemer in 2005, “*the resulting redundancy of cell death mechanisms has pathophysiological implications*”^10^. Cell death is inevitable and can be either physiological (e.g. for removing unwanted cells, such as cancer cells) or pathological (e.g. in neurodegenerative disorders). Nowadays, researchers noticed that there are at least 12 distinct types of RCD including various forms of regulated necrosis^7^. The term “regulated” indicates that RCD relies on fine-tuned molecular machinery involving signaling cascades and effectors. Thus, it is possible to find a way to characterize small chemical compounds that can modulate these pathways (such as with necrostatins for necroptosis^11^). These results led to the hypothesis that RCDs are « druggable », an emerging breakthrough that carries the potential to find new therapeutic approaches for unmet therapeutic needs including inflammatory and neurodegenerative diseases^12^. Necroptosis, is a regulated cell necrosis route that can be activated under apoptosis-deficient conditions. Necroptosis depends notably on the serine/threonine kinase activity of RIPK1 (Receptor-Interacting Protein Kinase 1) and RIPK3 and on the trafficking and accumulation at the plasma membrane of the pseudokinase MLKL (Mixed lineage kinase domain-like)^12,13^. As of December 2020, four RIPK1 inhibitors have reached clinical trials (GSK2982772 and GSK3145095 developed by GSK and DNL747 and DNL758 developed by Sanofi and Denali Therapeutics) for treatments of: psoriasis (phase II), ulcerative colitis (phase II), rheumatoid arthritis (phase II) or pancreatic cancer (phase II) for the molecules developed by GSK; Alzheimer’s disease (phase I), amyotrophic lateral sclerosis and multiple sclerosis (phase I) and for treatment of adult patients hospitalized with severe coronavirus disease 2019 (COVID-19) for the molecules developed by Sanofi and Denali Therapeutics (data from clinicaltrials.gov and^14^). Numerous other compounds (40+) were described having various binding mode on RIPK1^14^.

We contributed to this intensive hunt for potent RIPK1 inhibitors with the characterization of synthetic and natural-derivatives compounds including the 7-azaindole derivative Sibiriline^15^, the marine-derivative 2-aminobenzothiazole MBM105^16^ and flavanone nigratine (also known as 6E11)^17^. Nigratine (2-(4-(benzyloxy)phenyl)-2,5-dihydroxy-7-methoxychroman-4-one) is a synthetic derivative of the naturally occurring 2,5-dihydroxy-2-phenylchroman-4-ones isolated from *Populus nigra* buds. We previously showed that nigratine is a non-ATP competitive inhibitor with a remarkable selectivity toward RIPK1 and able to protect human aortic endothelial cells (HAEC) from cold hypoxia/reoxygenation-induced cell death^17^.

We now reported here the characterization of nigratine as new inhibitor of ferroptosis cell-death (with EC_50_ in the µM range). Nigratine is thus considered as the first-in-class dual inhibitor (or “magic shotgun”) of both necroptosis and ferroptosis regulated cell death that can be used in polypharmacological approaches for treatments of regulated-necrosis related disorders.

## RESULTS

### Effect of nigratine on necroptotic cell-death induced by TNF-α

Nigratine was characterized by Delehouzé et al. as new necroptosis inhibitor using a TNF-induced FADD-deficient human Jurkat necroptosis assay^17^. TNF-α can induce necroptosis in Jurkat cells when FADD is deleted. MTS assay was used to measure the viability of lymphocytes and the effect on the RIPK1-kinase activity was evaluated to study the mechanism of action of nigratine. In this study, we compared the bioactivity of nigratine with necrostatin-1 (nec-1) as control inhibitor of necroptosis^11^. These compounds were tested at the concentration of 10 and 50µM. The results obtained and the chemical structures of these compounds are depicted in Figure 1. In this assay, nigratine inhibited both necroptosis cell death and RIPK1 kinase activity with a similar activity than nec-1 (with both EC_50_ and IC_50_ in the μM range). The value of the protection index (dubbed ProtecΔ) for nigratine at 10µM (45.0) compared to the value at 50µM (26.0) suggested a lesser effect of the molecule on Jurkat FADD-deficient cells at high concentration.

**Figure 1:**
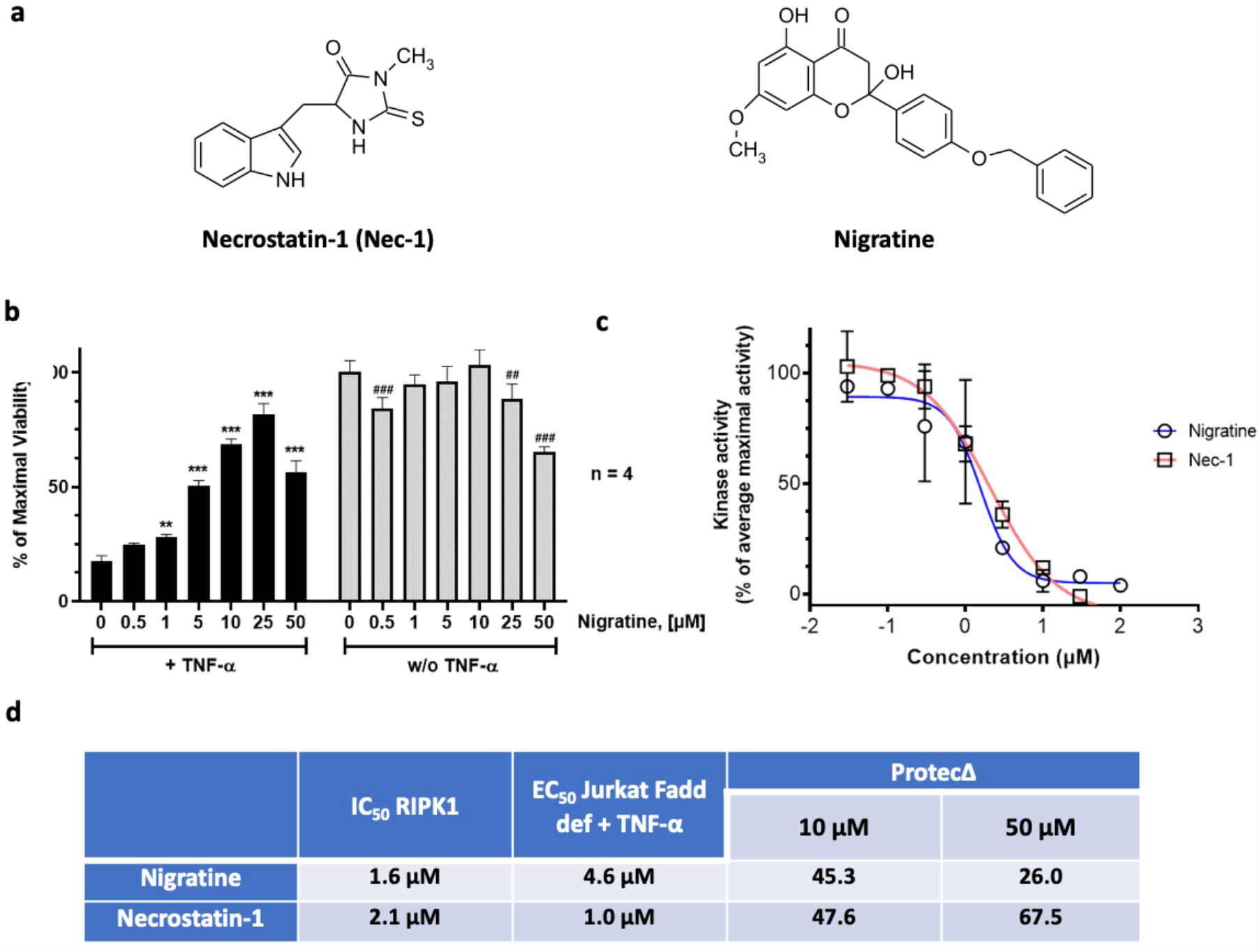
Nigratine inhibits necroptosis by targeting RIPK1 kinase. (**a**) Nigratine and necrostatin-1 (Nec-1), as control inhibitor of necroptosis, are structurally non related molecules. (**b**) Jurkat-Fadd def cells were treated 24h with increasing concentrations of nigratine and necrostatin-1 (Nec-1) alone or co-treated with 10ng/mL of TNF-α. Cell viability was estimated by MTS assay. Data are shown as the mean ± SD of four replicates. (**c**) The RIPK1 catalytic activity was monitored using myelin basic protein, MBP, as a substrate. RIPK1 was assayed in the presence of increasing concentrations of both nigratine and Nec-1. The kinase activities are expressed in % of average maximal activity, i.e. measured in the presence of DMSO (n=2, mean ± SEM). (**d**) The IC_50_ (on kinase) and EC_50_ and “protection delta” (ProtecΔ) (on cells) values were determined from the dose-response curves using Graphpad PRISM software and reported in the table. The level of protection against the TNF-α-induced cell death is estimated using the ProtecΔ values (first described and used in^16^). These data are determined by subtracting the value obtained for cells only treated with TNF-α from the value obtained for co-treatment with TNF-α and with the tested compound.

### Nigratine protects neuronal cells against ferroptotic cell death

In Delehouzé et al., we previously showed that treatment of human aortic endothelial cells (HAEC) with nigratine during cold hypoxia or during cold hypoxia and reoxygenation brought measurable benefits on cell survival. Compared to nec-1s, as specific RIPK1 inhibitor of necroptosis, the effect of nigratine was significantly better^17^. We thus suggested that this observed effect of nigratine was not fully related to the inhibition of RIPK1 kinase. Indeed, when compared to nigratine, nec-1s is a more potent inhibitor of RIPK1^17^.

We next analyzed the effect of nigratine treatment on ferroptotic cell-based models. Ferroptosis was first characterized in *NRAS*-mutant HT-1080 fibrosarcoma cells treated with erastin^8^. Erastin, like excess of L-glutamate (5mM), are class I ferroptosis inducers. They inhibit cystine uptake by the heterodimeric cystine/glutamate antiporter (system xc−) inducing an oxidative toxicity. This antiporter is thus a key component essential for protection of cells from oxidative stress and lethal lipid peroxidation, major hallmarks of ferroptosis. Glutathione peroxidase (GPX4) is also a key player of ferroptosis. The execution of ferroptosis depends on massive lipid peroxidation, which could be reduced by the lipid repair enzyme GPX4. RSL3 belongs to class II of ferroptosis-inducing agents by inhibiting GPX4^18^.

We tested here the protective effect of nigratine against class I (5mM glutamate and erastin) and II (RSL3) ferroptosis inducers in two neuronal cell lines, mouse HT22 (hippocampal neuronal cell line) and human SH-SY5Y (neuroblastoma cell line). These cell lines were widely used as models in studies of neurodegenerative processes, neurotoxicity, and neuroprotection. As shown on Figures 2 and 3, nigratine at 50µM was able to strongly inhibit the ferroptotic cell death induced by excess of glutamate, erastin and RSL3 and in both models of murine HT22 and human SH-SY5Y cell lines.

**Figure 2:**
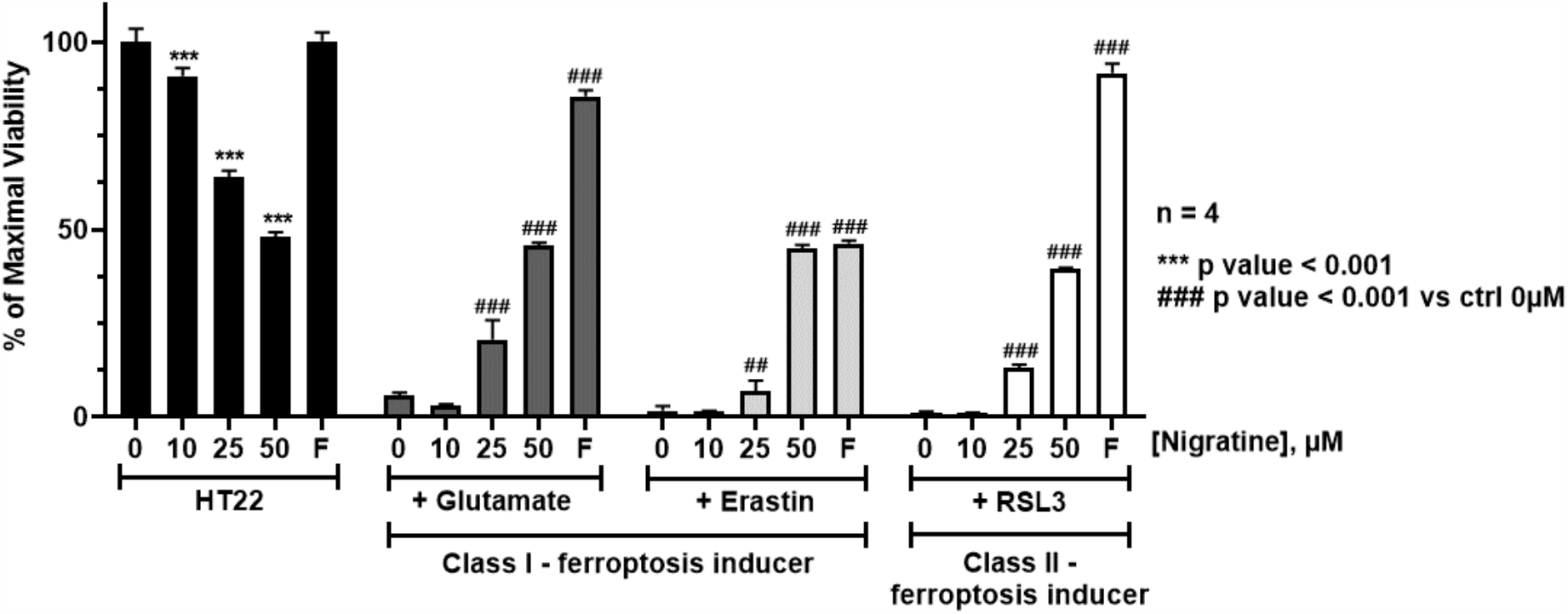
Nigratine protects HT-22 mouse hippocampal neuronal cell line from cell death triggered by both class I and II-inducers of ferroptosis. HT22 cells were treated 24h with increasing concentrations (0 - 50µM) of the tested chemical compounds alone (“HT22”) or co-treated with 5mM glutamate, 0.5µM erastin or 1µM of RSL3. Ferrostatin-1 (“F”) was used as a positive control for ferroptosis inhibition. Cell viability was estimated by MTS assay. Data are shown as the mean ± SEM of two replicates.

**Figure 3:**
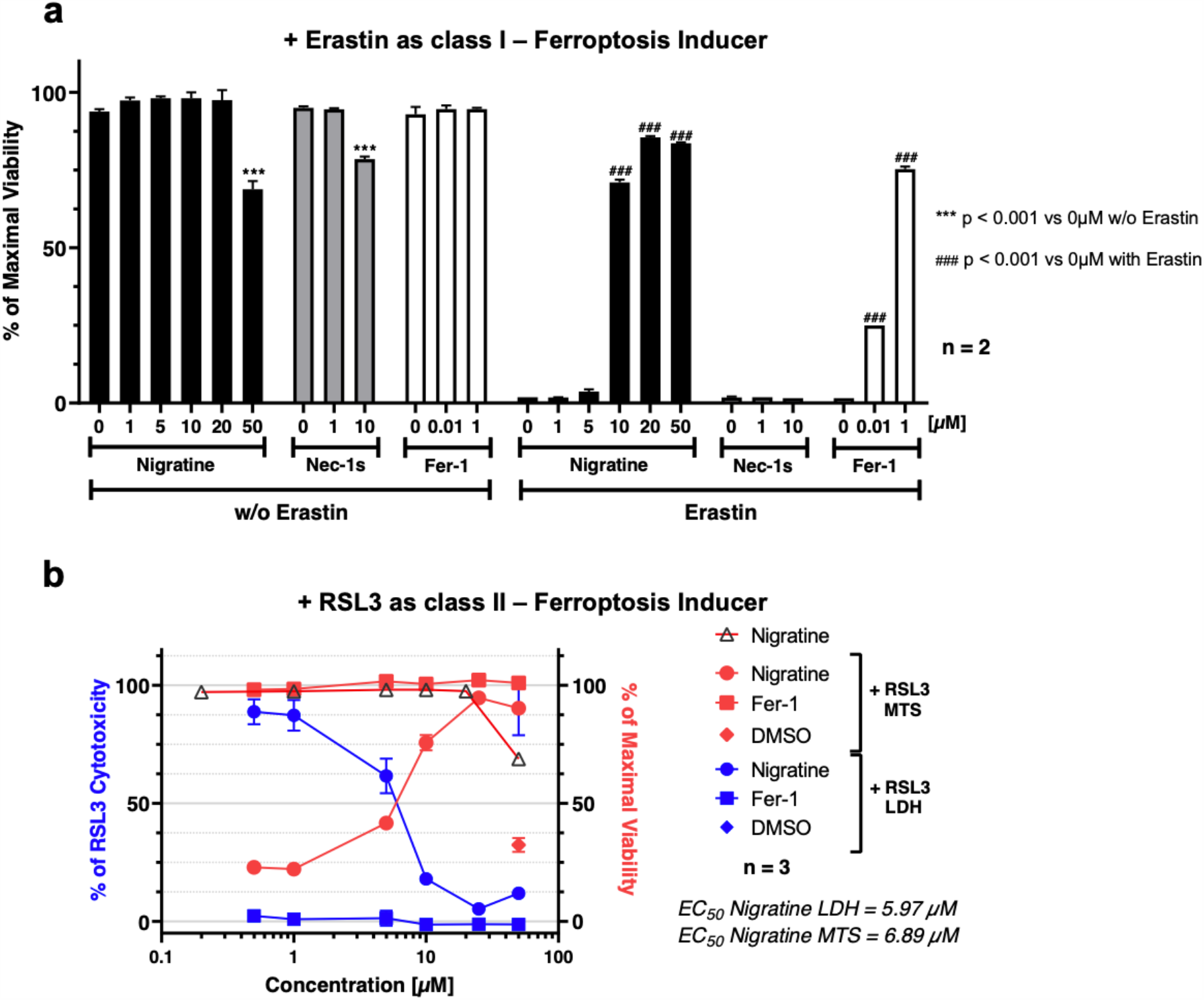
Nigratine protects SH-SY5Y human neuroblastoma cell line from cell death triggered by both class I and II-inducers of ferroptosis. (**a**) SH-SY5Y cells were co-treated 24h with various concentrations of the tested chemical compounds alone and 10µM of erastin. Ferrostatin-1 (Fer-1) was used as a positive control for ferroptosis inhibition. Cell viability was estimated by MTS assay. Data are shown as the mean ± SEM of two replicates. (**b**) SH-SY5Y cells were treated 24h with 5µM of RSL3 and increasing concentrations of nigratine or ferrostatin-1 (Fer-1). Cell death was evaluated by lactate dehydrogenase (LDH) release assay. Results are plotted in % of LDH release measured when cells are treated with RSL3 (left axis, colored blue). Cell viability was measured by MTS reduction assay. The results obtained (colored red) were plotted in % of maximal viability (detected in DMSO-treated cells, right axis). Data are shown as the mean ± SD of three replicates.

The dose-dependent effect of nigratine on cell viability was analyzed using MTS reduction assay. In order to validate the effect of nigratine on neuroprotection and necrosis, the extracellular lactate dehydrogenase (LDH) detection assay was also used as an independent cell death assay^19^. Indeed, the LDH is released into extracellular space when the plasma membrane is damaged. The necrotic cell death is essentially associated with the membrane permeabilization resulting to the rapid release of the cellular contents (including LDH and various damage-associated molecular patterns)^20^. As shown in Figure 3b, the treatment of SH-SY5Y with increasing doses of nigratine protected cells against cell-death in a dose dependent manner that correlated with the similar dose-dependent increase in cell survival. The results obtained also showed that nigratine has an acceptable cytotoxic effect on SH-SY5Y neuroblastoma cell line (Figure 3b). Interestingly, nigratine is active against ferroptosis on SH-SY5Y with EC_50_ around 6.5µM (EC_50(LDH)_=6.0µM and EC_50(MTS)_=6.9µM) in the same range than the activity against necroptosis (EC_50(MTS)_=4.6µM). Ferrostatin-1 was used as model inhibitor of ferroptosis and showed a better cellular effect compared to nigratine (Figure 2 and 3).

### Nigratine inhibits phospholipid peroxidation

We next selected the porcine renal epithelial cell line (LLC-PK1), a well-characterized renal proximal tubule cell line, to characterize the effect of nigratine on the peroxidation of phospholipids induced by the inhibition of GPX4 with RSL3. The results obtained are reported on Figure 4, indicated that: (i) nigratine protects renal epithelial cells against ferroptotic cell death induced by RSL3 (Figure 4a) and (ii) nigratine inhibits significantly phospholipid peroxide formation detected in cells using BODIPY 581/591 C11 probe (Figure 4b-c). This dye is a well-known sensitive indicator of free radical processes that have the potential to oxidize lipids in membranes. Taken together, these results suggested that nigratine has an effect on lipid peroxydation, one major hallmark of ferroptosis.

**Figure 4:**
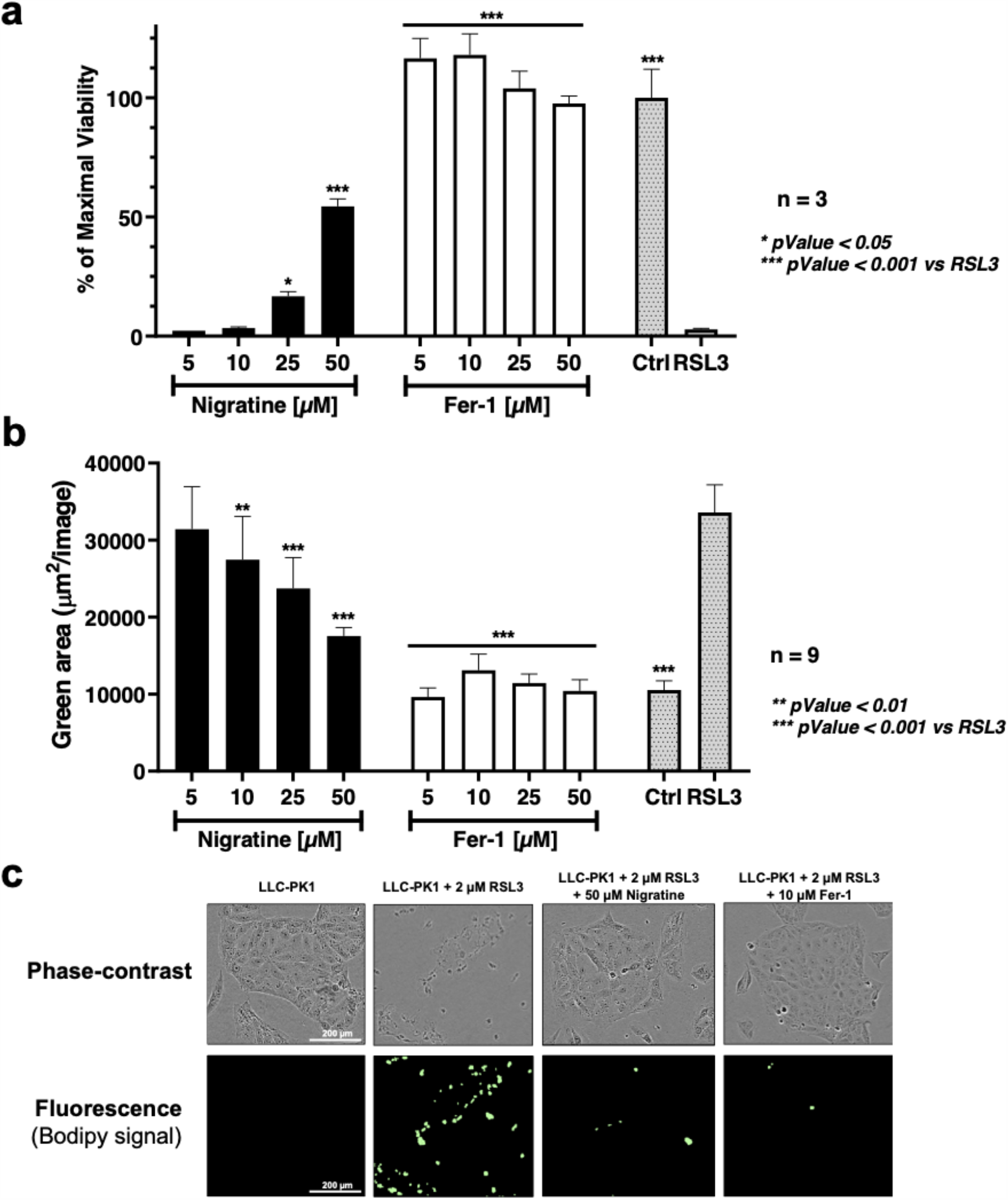
Nigratine protects porcine kidney epithelial LLC-PK1 cell line from lipid peroxidation and cell death triggered by RSL3. LLC-PK1 cell line was treated with 2µM of RSL3 and increasing concentration of Nigratine or ferrostatin-1 (Fer-1). (**a**) Cell viability was estimated by MTS assay. Data are shown as the mean ± SD of three replicates. (**b**) Lipid peroxidation was detected by cellular BODIPY 581/591 C11 staining. Fluorescence was recorded with the IncuCyte S3 live cell imaging apparatus. Data are shown as the mean ± SD of three replicates of nine replicates. (**c**) Representative phase-contrast and fluorescence images of cells stained with BODIPY 581/591 C11 probe are visualized using the IncuCyte S3 live cell imaging apparatus

### Nigratine is a weak antioxidant compound

Ferroptosis is characterized as a consequence of defects in antioxidant defenses. Indeed, ferroptosis is catalyzed by iron and is due to a loss of activity of the lipid repair enzyme glutathione peroxidase 4 (GPX4)^21^. The failure of the glutathione dependent antioxidant defenses leads to an accumulation of lipid-based reactive oxygen species (ROS), resulting of lipids peroxidation. Antioxidant compounds such as α-tocopherol (vitamin E) were shown to inhibit ferroptotic cell-death. α-tocopherol is a potent radical-trapping antioxidant (RTA) that breaks the auto-oxidation of chain-propagating peroxyl radicals and protects hydrocarbon biological systems from oxidation and membrane damage in ferroptosis (see^22^ for review). The results obtained (Figure 5) suggested that nigratine is only a weak antioxidant compound compared to α-tocopherol and the lipophilic antioxidant ferrostatin-1.

**Figure 5:**
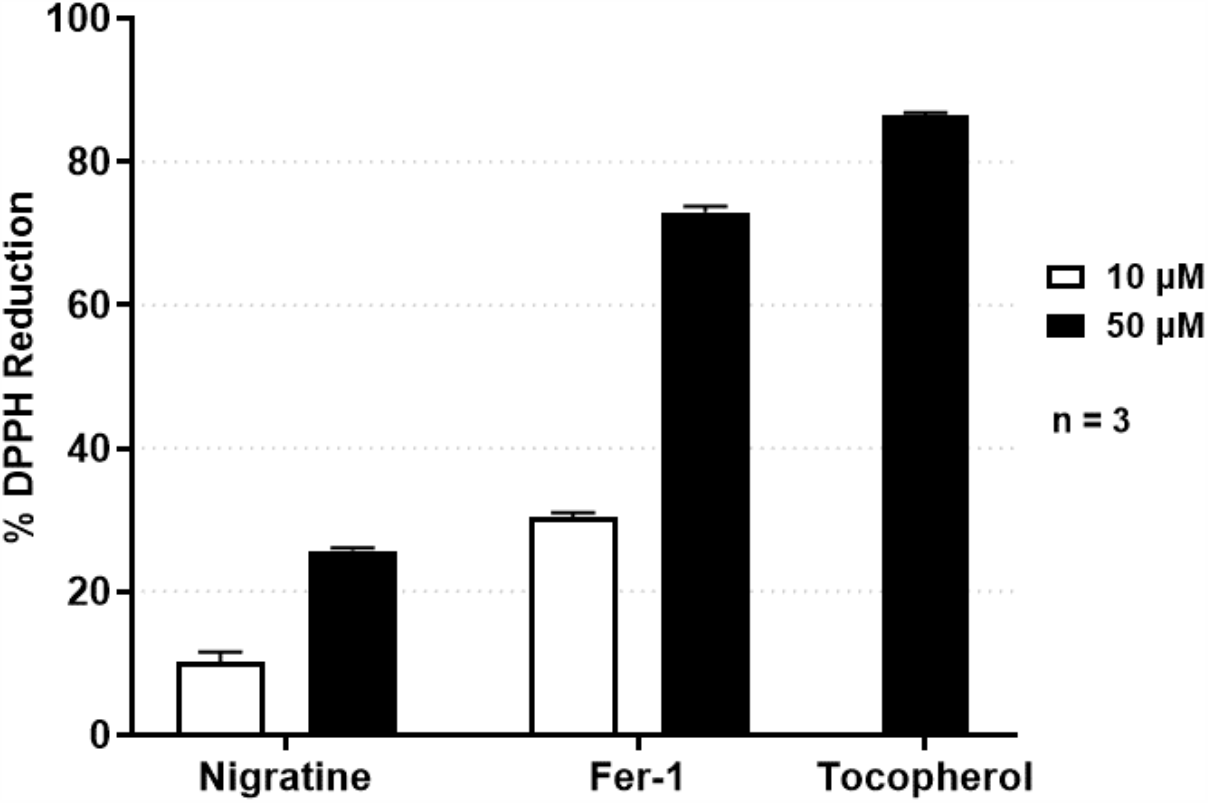
Antioxidant activity of nigratine. DPPH reduction assay was conducted for 30 min at room temperature after the addition of a DPPH solution at 15,85µM (in 90/10, vol/vol, methanol/water) in 96-well plate. Reduction of DPPH was determined by absorbance measurement at 517nm. Tocopherol (vitamin E), a well-known antioxidant, and Fer-1, as lipophylic antioxidant, were used as a positive controls. Data are shown as the mean ± SD of three replicates.

## DISCUSSION

Nigratine was previously described as a putative non-ATP competitive type III RIPK1 kinase inhibitor that can block the necroptosis cell death on various cellular models including Jurkat lymphocytes with EC_50_ in the µM range^17^. Necroptosis, a programmed cell death route, is clearly distinct from apoptosis as it does not involve key apoptosis regulators, such as caspases, Bcl-2 family members or cytochrome c release from mitochondria. In the present study, we showed that nigratine is also an inhibitor of ferroptosis: nigratine suppressed cell death induced by class I and II ferroptosis inducers: excess of glutamate, erastin and RSL3, respectively. We also showed that nigratine inhibits phospholipid peroxidation induced by RSL3 in epithelial LLC-PK1 porcine renal cells, an inhibition that was also observed for ferrostatin-1 as control molecule.

As mentioned here before, polypharmacological approaches -using drugs acting on multiple targets-appear suited to improve the outcome of complex diseases. Thus, nigratine appears to be very attractive in therapy for preventing and/or treating disorders associated with the induction of both necroptosis and ferroptosis. As example, the therapeutic benefit of nigratine can be evaluated in animal models of cisplatin-induced nephrotoxic acute kidney injury (AKI) where these two regulated necrosis were shown to be involved in proximal tubular cell death^23-25^. Ischemia/reperfusion injuries (IRI), that have been attributed to cell necrosis for decades^26^, should also be explored with nigratine.

Canonical ferroptosis inhibitors like the lipophilic antioxidant ferrostatin-1, prevent the accumulation of toxic lipid and cytosolic ROS, inhibiting ferroptotic cell death. As observed here using the DPPH assay, nigratine is comparatively less potent antioxidant than the two aromatic amines ferrostatin-1 and α-tocopherol. We consequently hypothesized that nigratine does not use a mechanism linked to the peroxyl radical scavenging for the inhibition of ferroptosis – or may inhibit another cellular target contributing to the overall cell-death protection. We now have to identify the molecular target of nigratine in ferroptosis using various approaches and notably by target fishing (see^27^ for review). This identification is prerequisite for improving the efficacy of dual-target inhibitors on ferroptosis-related phenotypes without affecting the activity of its primary target, the receptor-interacting protein kinase 1 (RIPK1). Numerous approaches are available to increase the efficacy of this class of drugs on rational basis. They include notably the combination of medicinal chemistry and computational strategies as it was already described for dual-target kinase inhibitors^28^.

Taken together these results obtained on nigratine indicate that pharmaceutical agents acting on both necroptosis and ferroptosis cell death routes can be designed and used to treat complex diseases involving the activation of multiple regulated necrosis. This study shed light on the emergence of polypharmacological approaches for treating multiple disorders where necrosis is of central pathophysiological relevance, such as: ischemia-reperfusion injury in brain, heart and kidney, inflammatory diseases, sepsis, retinal disorders or neurodegenerative diseases.

## METHODS

### Cell lines and culture

SH-SY5Y and HT22 cell lines were obtained from American Type Culture Collection (ATCC, Manassas, VA, USA) and were maintained in Dulbecco’s modified eagle’s medium (DMEM) containing 10% fetal bovine serum (FBS) in a humidified atmosphere of 5% CO_2_ at 37°C. LLC-PK1 was obtained from the European Collection of Authenticated Cell Cultures (ECACC, Porton Down, Salisbury, UK) and maintained in M199 medium supplemented with 10% FBS and cultured in humidified atmosphere at 37°C under 5% CO_2_. Medium and serum were purchased from Thermo Fisher Scientific (Gibco, Waltham, MA, USA).

### Reagents

Necrostatin-1 (Nec-1) and necrostatin-1s (Nec-1s) were from Calbiochem (VWR International, Fontenay-sous-Bois, FR), Ferrostatin-1 (Fer-1), Ras-selective lethal small molecule (RSL3) were from Selleckchem (Houston, TX, USA), 2,2-Diphenyl-1-picrylhydrazyl (DPPH) and α-tocopherol (vitamin E) was from Sigma Aldrich (St. Louis, MO, USA). TNF-α was obtained from Invitrogen (Carlsbad, CA, USA). This cytokine was used in the cell-based assay for the characterization of necroptosis inhibitors. Nigratine was obtained from the ICBMS institute (UMR5246, CNRS and University Claude Bernard Lyon 1, Lyon, France). The chemical synthesis of nigratine was described by Hauteville *et al*^*29*^.

### Cell death and cell viability assays

SH-SY5Y and HT22 cells were seeded in 96-well plates at a density of 10 000 or 5000 cells per well respectively following overnight incubation. Cells were treated with 5µM (SH-SY5Y cells) or 1µM (HT22 cells) of RSL3; 10µM (SH-SY5Ycells) or 0.5µM (HT22 cells) of erastin or 5mM (HT22 cells) of glutamate and increasing concentrations of compounds for 24h (100µl per well).

Cell death was determined by measurement of lactate dehydrogenase (LDH) leakage using the LDH Cytotoxicity assay kit (Invitrogen, Carlsbad, CA, USA) following manufacturer’s recommendations. LDH is a cytosolic enzyme that is rapidly released into the supernatant after cell damage. After 24h of treatment 50µl of supernatants were transferred into a clean 96-well plates, reagents were added and LDH activity was measured using a microplate reader. Percentage of cytotoxicity was calculated by dividing the LDH activity of the compounds with RSL3 by that of the RSL3 and DMSO.

Cell viability was assessed by MTS assay (CellTiter 96^®^ AQueous Non-Radioactive Cell Proliferation Assay; Promega, Fitchburg, WI, USA) according to the manufacturer’s instructions. This assay is based in the reduction of the 3-(4,5-dimethylthiazol-2-yl)-5-(3-carboxymethoxyphenyl)-2-(4-sulfophenyl)-2H-tetrazolium (MTS) by viable cells to form a colored formazan product. After treatment, cells were incubated 3h at 37°C, 5% CO_2_ with MTS. The absorbance was measured using a microplate reader at 490 and 630nm and the percentage of viability was calculated by dividing the absorbance of testing compound by the absorbance of DMSO treated cells (control).

### RIPK1 kinase assay

Human RIPK1 full length GST-tagged was purchased from SignalChem (Richmond, CA, USA). The protocol used to detect the enzymatic activity was described in Delehouzé *et al*^17^. Briefly, RIPK1 kinase was performed on KISSf screening facility (IBISA, Biogenouest, Station Biologique, Roscoff, France) with myelin basic protein (MBP, Sigma, #M1891) as substrate and in the presence of 15µM ATP.

### Cell-based lipid peroxidation assay

Ferroptosis was induced by treatment of porcine LLC-PK1 cells (derived from the renal epithelial cells of Hampshire pigs PK1) with 2µM of RSL3. For lipid peroxidation assay, cells were seeded in 96-well black plates with clear bottom at a density of 10,000 cells per well. The assay was performed as previously described by Kahn-Kirby et al^30^. Briefly, after overnight incubation, cells were pre-labeled with 10µM of BODIPY 581/591 C11 dye (Invitrogen, Carlsbad, CA, USA) for 30min at 37°C; 5% CO_2_. Cells were washed 3 times with PSB before adding treatment. Cells were then treated with RSL3 (2µM) and increasing concentrations of nigratine or ferrostatin-1 (Fer-1) and incubated at 37°C with 5% CO_2_ during 24h in the IncuCyte S3 live-cell imaging and analysis system (Essen BioScience, Sartorius, Göttingen, Germany). Images were acquired using a 10X objective and 440-480 nm Excitation / 504-544 nm Emission filters, once hourly for up to 24h.

### DPPH reduction assay

Antioxidant activity of compounds was determined by a DPPH (2,2-Diphenyl-1-picrylhydrazyl) reduction assay. DPPH was prepared at 15.85µM in 90% methanol and added to several concentrations of compounds in 96-well plate (200µL per well). Tocopherol, a well-known antioxidant, was used as control. The reaction was conducted at room temperature for 30 minutes and absorbance was measured at 517nm using the EnVision microplate reader (PerkinElmer, Waltham, MA, USA). The percentage of DPPH reduction was calculated by dividing the difference between the absorbance of DPPH and those of compounds by the absorbance of DPPH.

### Statistical analyses

Data from a minimum of two experiments were expressed as means ± range; ± SD or ± SEM. Statistical analyses were done by ANOVA, Tukey’s Multiple Comparison Test and Student’s *t*-test for two groups of data, and significance levels used are \**P*<0.05, \*\**P* <0.01, \*\*\**P*<0.001 by using GraphPad Prism6 software (GraphPad Software, San Diego, CA, USA).

## ACKNOWLEDGMENTS

We thank Amandine Bescond for advices on DPPH assay and Blandine Baratte for RIPK1 inhibition assays. The authors also thank the GIS IBiSA (Infrastructures en Biologie Santé et Agronomie, France) and Biogenouest (Western France life science and environment core facilty network) for supporting KISSf screening facility (Roscoff, France). Claire Delehouzé is recipient of a CIFRE PhD fellowship. Stéphane Bach and Marie-Thérèse Dimanche-Boitrel were supported by SATT Ouest Valorisation (“Gref-Pres” maturation program). Stéphane Bach is supported by the Fondation d’Entreprise Grand Ouest (“Houarnine” program). SeaBeLife is supported by the Technopole Brest Iroise, BPI, Biotech Santé Bretagne and Région Bretagne.

## AUTHOR CONTRIBUTIONS

CD designed, performed the cell-based screening of the chemical library, performed and analyzed *in vitro* data and co-wrote the manuscript. AC and MH synthesized the compounds and characterized them by RMN and mass spectrometry. PG and MTDB analyzed results of chemical synthesis and biological evaluation, respectively, and revised the manuscript. MR and SB supervised the study (design and coordination) and were the co-promotor of the PhD of CD. MR revised the manuscript. SB wrote the manuscript. All the authors read and approved the final manuscript submitted for publication.

## REFERENCES

1 Gandini, A. et al. Tau-Centric Multitarget Approach for Alzheimer’s Disease: Development of First-in-Class Dual Glycogen Synthase Kinase 3beta and Tau-Aggregation Inhibitors. J Med Chem 61, 7640–7656, doi:10.1021/acs.jmedchem.8b00610 (2018).

2 Albertini, C., Salerno, A., Sena Murteira Pinheiro, P. & Bolognesi, M. L. From combinations to multitarget-directed ligands: A continuum in Alzheimer’s disease polypharmacology. Medicinal Research Reviews, doi:10.1002/med.21699 (2020).

3 Ehrlich, P. Beitråge zur Theorie und Praxis der histologischen Fårbung. The Collected Papers of Paul Ehrlich, London Pergamon, 29–64 (2013).

4 Roth, B. L., Sheffler, D. J. & Kroeze, W. K. Magic shotguns versus magic bullets: selectively non-selective drugs for mood disorders and schizophrenia. Nat Rev Drug Discov 3, 353–359, doi:10.1038/nrd1346 (2004).

5 Rosini, M. Polypharmacology: the rise of multitarget drugs over combination therapies. Future Medicinal Chemistry 6, 485–487, doi:10.4155/fmc.14.25 (2014).

6 Ramsay, R. R., Popovic-Nikolic, M. R., Nikolic, K., Uliassi, E. & Bolognesi, M. L. A perspective on multi-target drug discovery and design for complex diseases. Clinical and Translational Medicine 7, pdoi:10.1186/s40169-017-0181-2 (2018).

7 Galluzzi, L. et al. Molecular mechanisms of cell death: recommendations of the Nomenclature Committee on Cell Death 2018. Cell death and differentiation 25, 486–541, doi:10.1038/s41418-017-0012-4 (2018).

8 Dixon, S. J. et al. Ferroptosis: an iron-dependent form of nonapoptotic cell death. Cell 149, 1060–1072, doi:10.1016/j.cell.2012.03.042 (2012).

9 Aguilera, A. et al., doi:10.1101/828293 (2019).

10 Golstein, P. & Kroemer, G. Redundant cell death mechanisms as relics and backups. Cell Death & Differentiation 12, 1490–1496, doi:10.1038/sj.cdd.4401607 (2005).

11 Degterev, A. et al. Chemical inhibitor of nonapoptotic cell death with therapeutic potential for ischemic brain injury. Nat Chem Biol 1, 112–119, doi:10.1038/nchembio711 (2005).

12 Degterev, A., Ofengeim, D. & Yuan, J. Targeting RIPK1 for the treatment of human diseases. Proceedings of the National Academy of Sciences 116, 9714–9722, doi:10.1073/pnas.1901179116 (2019).

13 Samson, A. L. et al. MLKL trafficking and accumulation at the plasma membrane control the kinetics and threshold for necroptosis. Nature Communications 11, pdoi:10.1038/s41467-020-16887-1 (2020).

14 Martens, S., Hofmans, S., Declercq, W., Augustyns, K. & Vandenabeele, P. Inhibitors Targeting RIPK1/RIPK3: Old and New Drugs. Trends in pharmacological sciences 41, 209–224, doi:10.1016/j.tips.2020.01.002 (2020).

15 Le Cann, F. et al. Sibiriline, a new small chemical inhibitor of receptor-interacting protein kinase 1, prevents immune-dependent hepatitis. FEBS J 284, 3050–3068, doi:10.1111/febs.14176 (2017).

16 Benchekroun, M. et al. Discovery of simplified benzazole fragments derived from the marine benzosceptrin B as necroptosis inhibitors involving the receptor interacting protein Kinase-1. European Journal of Medicinal Chemistry 201, pdoi:10.1016/j.ejmech.2020.112337 (2020).

17 Delehouze, C. et al. 6E11, a highly selective inhibitor of Receptor-Interacting Protein Kinase 1, protects cells against cold hypoxia-reoxygenation injury. Sci Rep 7, 12931, doi:10.1038/s41598-017-12788-4 (2017).

18 Conrad, M., Angeli, J. P., Vandenabeele, P. & Stockwell, B. R. Regulated necrosis: disease relevance and therapeutic opportunities. Nat Rev Drug Discov 15, 348–366, doi:10.1038/nrd.2015.6 (2016).

19 Chan, F. K.-M., Moriwaki, K. & De Rosa, M. J. in Immune Homeostasis Methods in Molecular Biology Ch. Chapter 7, 65–70 (2013).

20 Grootjans, S., Vanden Berghe, T. & Vandenabeele, P. Initiation and execution mechanisms of necroptosis: an overview. Cell Death & Differentiation 24, 1184–1195, doi:10.1038/cdd.2017.65 (2017).

21 Yang, W. S. & Stockwell, B. R. Ferroptosis: Death by Lipid Peroxidation. Trends in cell biology 26, 165–176, doi:10.1016/j.tcb.2015.10.014 (2016).

22 Kajarabille& Latunde, D. Programmed Cell-Death by Ferroptosis: Antioxidants as Mitigators. International Journal of Molecular Sciences 20, pdoi:10.3390/ijms20194968 (2019).

23 Hu, Z. X. et al. VDR activation attenuate cisplatin induced AKI by inhibiting ferroptosis. Cell death & disease 11, pdoi:10.1038/s41419-020-2256-z (2020).

24 Deng, F., Sharma, I., Dai, Y. B., Yang, M. & Kanwar, Y. S. Myo-inositol oxygenase expression profile modulates pathogenic ferroptosis in the renal proximal tubule. Journal of Clinical Investigation 129, 5033–5049, doi:10.1172/Jci129903 (2019).

25 Xu, Y. F. et al. A Role for Tubular Necroptosis in Cisplatin-Induced AKI. Journal of the American Society of Nephrology 26, 2647–2658, doi:10.1681/Asn.2014080741 (2015).

26 Brady, H. R. & Singer, G. G. Acute renal failure. The Lancet 346, 1533–1540, doi:10.1016/s0140-6736(95)92057-9 (1995).

27 Guiffant, D. et al. Identification of intracellular targets of small molecular weight chemical compounds using affinity chromatography. Biotechnol J 2, 68–75 (2007).

28 Sun, D. et al. Dual-target kinase drug design: Current strategies and future directions in cancer therapy. European Journal of Medicinal Chemistry 188, pdoi:10.1016/j.ejmech.2019.112025 (2020).

29 Hauteville, M., Chopin, J., Geiger, H. & Schuler, L. Protogenkwanin, a New Flavonoid from Equisetum-Arvense L. Tetrahedron 37, 377–381, doi:Doi 10.1016/S0040-4020(01)92024-1 (1981).

30 Kahn-Kirby, A. H. et al. Targeting ferroptosis: A novel therapeutic strategy for the treatment of mitochondrial disease-related epilepsy. Plos One 14, pdoi:10.1371/journal.pone.0214250 (2019).

